# Cryptic recombination and transposition drive structural variation to shape genomic plasticity and life history traits in a host generalist fungal plant pathogen

**DOI:** 10.1101/2024.07.02.600549

**Authors:** Mark C Derbyshire, Toby E Newman, Yuphin Khentry, Pippa J Michael, Sarita Jane Bennett, Ashmita Rijal Lamichhane, Carolyn Graham-Taylor, Subhash Chander, Claudia Camplone, Simone Vicini, Laura Esquivel-Garcia, Cathy Coutu, Dwayne Hegedus, John Clarkson, Kurt Lindbeck, Lars G Kamphuis

## Abstract

**Background:** An understanding of plant pathogen evolution is important for sustainable management of crop diseases. Plant pathogen populations must maintain adequate heritable phenotypic variability to survive. Polymorphisms >= 50 bp, known as structural variants (SVs), could contribute strongly to this variability by disrupting gene activities. SV acquisition is largely driven by mobile genetic elements called transposons, though a less appreciated source of SVs is erroneous meiotic double-strand break repair. The relative impacts of transposons and recombination on SV diversity and the overall contribution of SVs to phenotypic variability is elusive, especially in host generalists.

**Results:** We use 25 high quality genomes to create a graphical pan-genome of the globally distributed host-generalist crop pathogen *Sclerotinia sclerotiorum*. Outcrossing and recombination rates in this self-fertile species have been debated. Using bisulfite sequencing, and short read data from 190 strains, we show that *S. sclerotiorum* has many hallmarks of eukaryotic meiosis, including recombination hot and cold spots, centromeric and genic recombination suppression, and rapid linkage disequilibrium decay. Using a new statistic that captures average pairwise structural variation, we show that recombination and transposons make distinct contributions to SV diversity. Furthermore, despite only 5 % of genes being dispensable, SVs often had a stronger impact than other variants across 14 life history traits measured in 103 distinct strains.

**Conclusion:** Transposons and recombination make distinct contributions to SV diversity in *S. sclerotiorum*. Despite limited gene content diversity, SVs may strongly impact phenotypic variability. This sheds light on the genomic forces shaping adaptive flexibility in host generalists.

## Background

An understanding of the evolutionary processes underpinning plant pathogen adaptation is crucial for developing better disease management strategies, such as resistant cultivars, prediction of epidemics and monitoring of fungicide resistance [1–3]. Population genetic approaches can be used to understand the evolutionary characteristics of plant pathogens [4,5], although their application has been limited in the past to variants that can be confidently genotyped using short reads. However, with the now widespread use of long read sequencing, more plant pathogen pan-genomes of increasing quality are becoming available for evolutionary studies [6–12].

Aside from simple genotypic variants, such as single nucleotide polymorphisms (SNPs) and small insertions/deletions (InDels), complete genomes assembled using long reads can be used to identify structural variants (SVs), which are generally defined as polymorphisms of more than or equal to 50 bp [13]. These can be confidently genotyped using long reads assemblies and incorporated into a data structure known as a pan-genome graph [14,15]. Based on this underlying representation of genomic variation, SVs can be genotyped in a broader set of individuals using short reads [16,17]. This approach also improves the accuracy of non-SV calls by improving short read placement and reducing reference bias [18].

In many species, this and similar techniques have revealed that previously invisible SVs are strongly linked with phenotypic variability. For instance, the tomato graph pan-genome showed that SVs are a major component of ‘missing heritability’ [19], explaining much of the phenotypic variance not captured by simpler variants called against a single reference. Furthermore, in the important fungal wheat pathogen *Zymoseptoria tritici*, SVs have been shown to make a substantial contribution to important life history traits, such as tolerance of fungicides [20]

Two key processes underpinning evolutionary adaptation are mutation, including *de novo* acquisition of SVs, and meiotic recombination. The traditional view is that mutation creates new alleles and meiotic recombination shuffles alleles to create new haplotypes [21]. Shuffling of alleles into novel haplotypes allows beneficial alleles to spread without the burden of linked deleterious alleles. Without meiotic exchange, populations are likely to gradually accumulate deleterious mutations that cannot be lost without also losing beneficial mutations, a process known as Muller’s ratchet [22–24].

Though evolutionary theory often ascribes distinct roles to mutation and meiotic recombination in creating and shuffling alleles, respectively, the two processes may not be completely orthogonal. Meiosis itself may be powerfully mutagenic, as it requires the induction of numerous double-strand DNA breaks. Through erroneous repair of these breaks, meiosis has been linked with exceptionally high *de novo* mutation rates [12,20,21]. All types of mutations can occur through faulty repair of double-strand breaks, although meiosis-induced double strand breaks may be particularly prone to creating new SVs [21]. In humans, for example, double-strand breaks induced by meiosis lead to a 400 to 1,000-fold increase in the rate of SV acquisition, and many SVs induced in recombination hotspots are pathogenic, highlighting the impact of meiotic mutagenesis on phenotype and human disease [12, 25]. Recently, a machine learning approach showed that multiple genomic features, including local recombination rate, were highly predictive of SVs induced in haploid offspring of crosses of *Z. tritici* [20]. In the plant pathogen *Fusarium graminearum*, local recombination rate was also shown to be associated with SVs across four high quality genomes [12].

In addition to meiosis, transposition is a highly potent instigator of structural variation in genomes. This occurs when active mobile elements called transposons duplicate or relocate themselves in the genome [26]. In addition, the repetitive nature of transposons can create SVs through pairing of distant genomic copies during DNA damage repair via the homologous recombination pathway [26]. Though transposons can be destabilising to genomes, occasionally they create beneficial mutations, which are an important source of adaptive evolution [27].

In plant pathogens, transposition is widely appreciated as one of the main driving forces of genomic plasticity. Though meiotic exchange has been linked with *de novo* acquisition of SVs in plant pathogenic fungi, the link between meiotic exchange and genome stability has not been widely explored in plant pathogen populations, and little is known about how meiosis and transposition interact to shape SV diversity. Despite several long reads pathogen pan-genomes, the overall contribution of SVs to variability in life history traits is also poorly understood.

To date, much of the research on the evolution of plant pathogen genomes has also been conducted on host specialists, which are under acute selective pressure to maintain virulence on a single species. In contrast, the fungus *Sclerotinia sclerotiorum* infects hundreds of plant species in at least 74 documented families [28]. Though its genome may harbour some polymorphic regions [29,30], in contrast to many host specialists, its predicted effectors are largely conserved [30] and several are likely compatible with diverse hosts [31,32]. This suggests that, like many niche-generalists, *S. sclerotiorum* has evolved an energetically-optimised and multifunctional genome, which facilitates its colonisation of diverse hosts [33–35].

Sporulation in *S. sclerotiorum* occurs through obligate sexual reproduction. However, since it is self-fertile (homothallic), sexual reproduction can create genotypically uniform progeny, allowing certain genotypes to persist for long periods of time as clones [36]. The extent to which *S. sclerotiorum* outcrosses to generate new diversity has been debated, with some suggesting homothallism promotes universal outcrossing [29,37–44] and others suggesting that outcrossing is extremely rare [45–47]. Consequently, the overall contribution of meiotic exchange to genome stability and evolution in this species is particularly poorly understood.

Here, we present a global graphical pan-genome of *S. sclerotiorum* and use 25 reference- quality genomes and 190 short reads samples to investigate species-wide SV diversity. To capture this diversity, we present a new statistic called ‘SVπ’, which describes the average number of SVs between all pairs of individuals. Using population genetics techniques, we establish *S. sclerotiorum* as an outcrossing species with many of the hallmarks of eukaryotic meiotic recombination, such as rapid linkage disequilibrium decay, suppressed recombination at centromeres, recombination hot and cold spots and enhanced recombination outside of coding sequences. We find that both recombination rate and transposable element content are independently positively correlated with total number of SVs and SVπ though not positively correlated with one another.

Overall, unlike that of most host specialists studied to date, we show that gene content in the *S. sclerotiorum* genome is largely stable, despite numerous small, unstable, repeat-rich, gene-sparse regions. SVs often had a stronger effect than other variants on 14 life history traits assessed across 103 strains, and we find that a 48 bp InDel is significantly associated with tolerance of the fungicide azoxystrobin. Overall, our data suggest that transposition and meiotic recombination make distinct contributions to SV diversity in *S. sclerotiorum*, and that SVs may be an important driver of phenotypic plasticity, despite the stability in gene content of the species. These insights shed new light on the genomic processes underpinning the evolution of host generalism in plant pathogens.

## Results and discussion

### Development of a *Sclerotinia sclerotiorum* graphical pan-genome

To create a high-quality set of *S. sclerotiorum* genomes for SV analysis, we generated Illumina-corrected Oxford Nanopore long reads assemblies of the genomes of 24 diverse strains from Australia (10 strains), Europe (5 strains) and Canada (9 strains). Overall, 23 of the strains had telomere-to-telomere assemblies for >= 10 of the 16 *S. sclerotiorum* chromosomes, and seven had telomere-to-telomere assemblies for >= 14 chromosomes. There were few gaps in assemblies on average, and 22 strains had gapless assemblies for >= 10 chromosomes; three of these strains had gapless assemblies for all 16 chromosomes. All these assemblies are comparable to the reference *S. sclerotiorum* genome [30], which has 14 telomere-to-telomere and 15 gapless chromosomes. BUSCO scores ranged from 98.9 to 99.2, with a median of 99.05 (Supplementary Table 1), confirming the completeness of these assemblies. These new assemblies are available in NCBI under BioProject PRJNA1112094.

To explore structural variation in *S. sclerotiorum* we constructed a pan-genome graph from the genomes of these 24 strains and the reference strain. In this graph, we identified 186,486 variants, including 154,892 SNPs, 5,877 multiple nucleotide polymorphisms (MNPs), 20,061 InDels and 5,556 SVs. There were 9,876 complex variants with more than one allele, including 2,892 (52 %) of the SVs.

To capture more genotypic diversity, we aligned Illumina short reads to the pan-genome graph from an additional 190 strains, 181 of which were sequenced in this study (available in NCBI under BioProject PRJNA1120954) (Supplementary Table 2). Overall, the genotypes of 3,741 of the SVs from the pan-genome graph were captured in this broader data set. This data set is the first graphical pan-genome of the important host generalist pathogen *S. sclerotiorum*. It includes 215 strains, with 152 from Australia, 17 from North America, 44 from Europe, and one each from South Africa and Morocco.

## *Sclerotinia sclerotiorum* undergoes cryptic recombination whilst maintaining clonal lineages across large temporal and spatial distances

*S. sclerotiorum* produces ascospores through sexual reproduction. As it is homothallic, ascospores may be genotypically identical, which leads to an effectively clonal mode of propagation. Clonality is evident in the detection of temporally or spatially distant genotypically nearly identical strains. We identified 120 clonal lineages (>= 98 % identical) among the 215 strains (Supplementary Figure 1). Clonality was most prevalent among the Australian strains, whereas European and North American strains were mostly genotypically distinct (Figure 1). This was expected because most of the European and North American strains were previously shown to be distinct lineages using markers [48,49], whereas 99 of the Australian strains were collected from five sites (two of which were in the same locality) in Western Australia with no prior genotyping [50,51].

**Figure 1.**
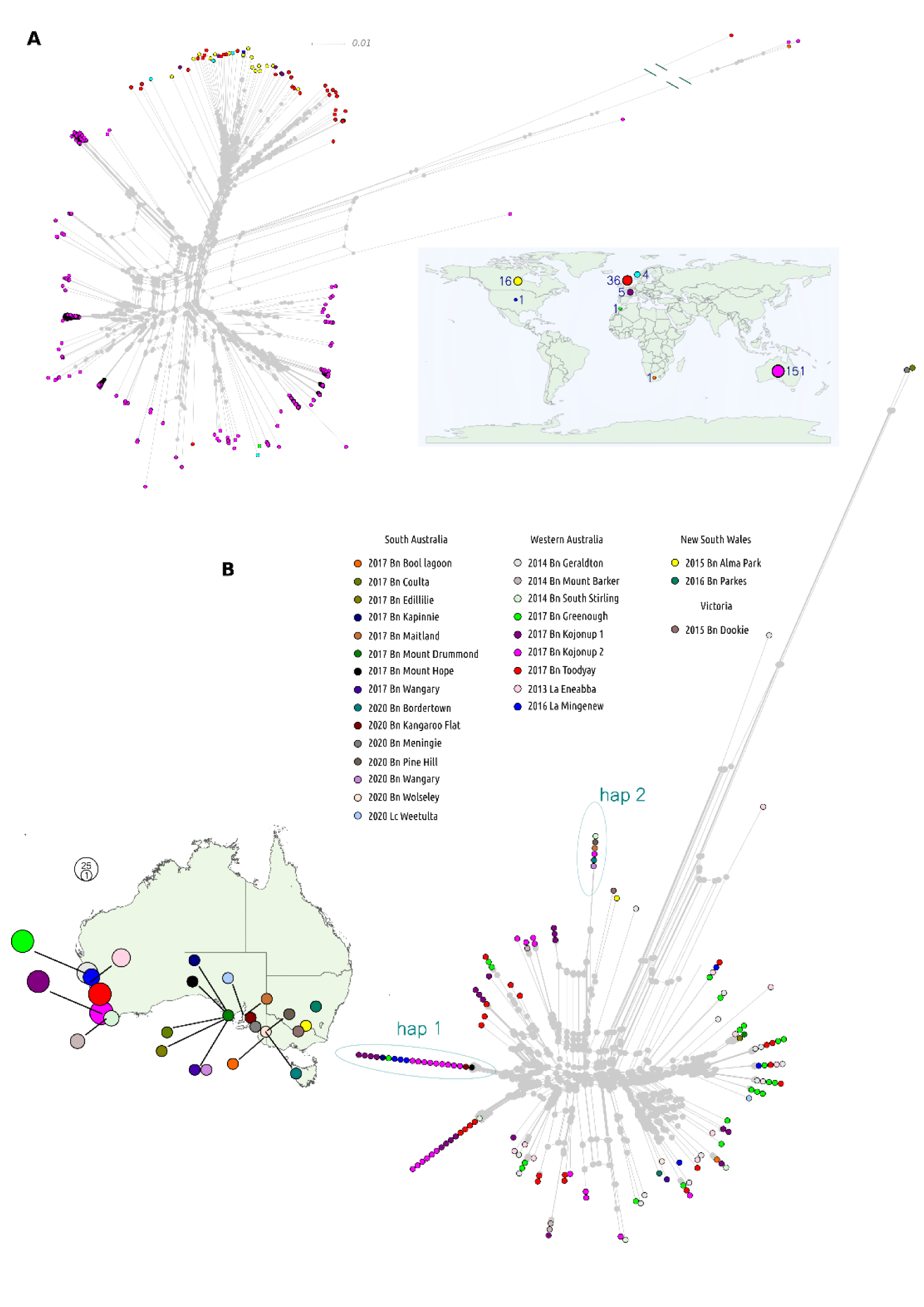
Genotypic clustering of *Sclerotinia sclerotiorum* strains from the global population sample. **A** A phylogenetic network with all strains in the dataset coloured according to geographical origin. The map inset shows where strains were collected with colours corresponding to those in the network. The sizes of circles on the map corresponds with the number of strains from each global region. **B** A phylogenetic network for the Australian strains. Circles are coloured according to geographical origin within Australia. Where circles are stacked on top of each other, isolates are a >= 98 % genotypically identical group of clones. The map to the left shows where isolates were collected within Australia, with colours of circles corresponding to colours on the network. The sizes of circles represent the numbers of strains from each collection site. Haplotypes 1 (hap 1) and 2 (hap 2) are examples of frequently-sampled and geographically-widespread clones, with individuals from Western Australia and South Australia.

Confirming the long-term maintenance of clonal propagation, we found several clones from geographically distant regions, some of which were collected many years apart. For example, strains CU11.18 and F19064 were found in Western Australia in 2013 and South Australia in 2018, respectively. The most extreme example was the pair of clones S55 and MB57, which were collected in the USA in 1987 and Manitoba in 2010, respectively.

In our previous study, we found that the global *S. sclerotiorum* population forms two distinct sub-populations, between which there has been limited gene flow. SNP data from the 215 genomes confirmed this observation (Figure 1), showing that Australian/African and European/North American strains formed mostly distinct sub-populations (referred to as AuAf and EuNA herein). Although we expanded the Australian collection, our study only contained the two African strains from our previous study [29], Sssaf from South Africa and Ss44 from Morocco, so the global relationship between Australian and African strains is still not fully resolved.

In the AuAf sub-population, we found evidence for three ancestral populations, and numerous admixed individuals. In the EuNA population, we identified three further ancestral populations with limited admixture (Figure 2 A). Two of the EuNA strains were admixed individuals containing alleles from either the AuAf ancestral populations or both the EuNA and AuAf ancestral populations. The widespread recent admixture among AuAf strains supports outcrossing between lineages from distinct ancestral populations.

**Figure 2.**
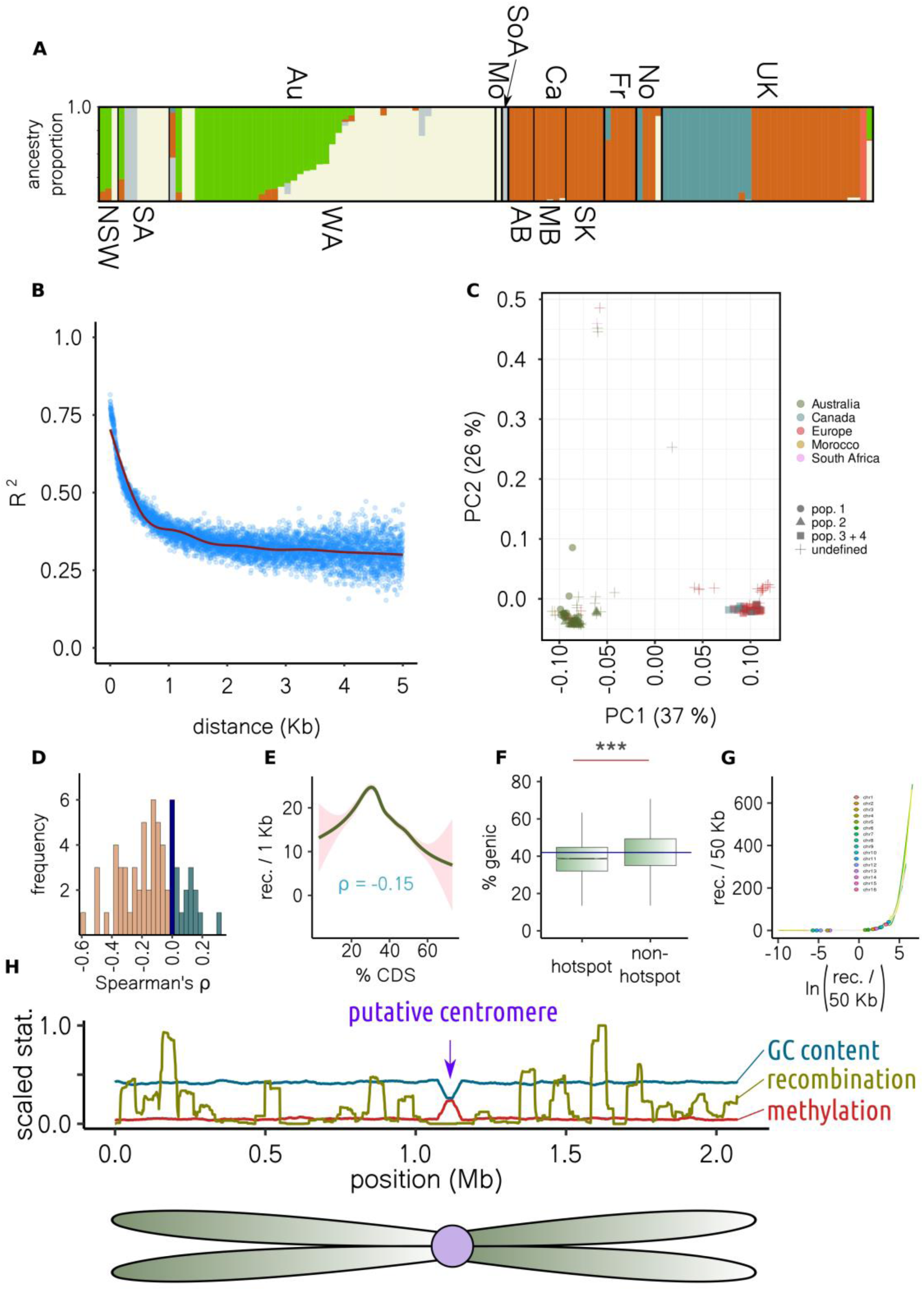
Population structure and evidence of recombination. **A** Colours correspond to ancestral populations making up individuals. Country of origin (above) is Au = Australia, Mo = Morocco, SoA = South Africa, Ca = Canada, Fr = France, No = Norway, and UK = UK. Below, states within Australia and Canada are indicated, where NSW = New South Wales, SA = South Australia, WA = Western Australia, AB = Alberta, MB = Manitoba, and SK = Saskatchewan. **B** Linkage disequilibrium (y axis) decay with physical distance (x axis). Points are averages for unique distance measurements, and the red line is a general additive model fit. **C** The first two principal components of genotypic variance. Colours indicate geographical origin and point shapes the four population sub-samples used for recombination analysis. **D** Across chromosomes and population sub-samples, the distribution of Spearman’s correlations between chromosome end distance and recombination rate. **E** Correlation between coding DNA sequence content (x axis) and recombination rate (y axis) of 50 Kb sliding windows. The line is a a general additive model fit. **F** Boxplot showing percent gene content of 50 Kb windows containing and not containing recombination hotspots (*** = P < 2e^-16^). Boxes and whiskers show interquartile range. **G** Circles show where windows containing putative centromeres lie on a plot of recombination rate (y axis) against log recombination rate (x axis). Putative centromeres are in regions of low recombination, before the inflection point. **H** The y axis is scaled (division by maximum) recombination rate, amount of methylation or GC content for sliding windows. The x axis shows position (Mb) across chromosome 6 (all chromosomes and population samples are in Supplementary File 1). All chromosomes had a dip in GC coincident with a spike in methylation, almost always coincident with a recombination cold spot.

To further explore outcrossing in the global *S. sclerotiorum* population, we investigated the rate of linkage disequilibrium decay in the 120 independent clonal lineages. We found that across the whole population, linkage disequilibrium decayed to half its maximum value at 428 bp (Figure 2 B). Three tests of the association between recombination and physical distance, neighbour similarity score [52], maximum Χ^2^ [53], and pairwise homoplasy index [54], also supported statistically significant recombination between non-adjacent alleles (P = 0 for all tests and chromosomes, Supplementary Table 3).

The rate at which LD decays to half its maximum (LD_2_) value is typically higher in predominantly outcrossing and lower in predominantly clonal species [55,56]. Species that rarely outcross often have LD_2_ rates of more than 100 Kb, whereas highly outcrossing species have rates on the order of a few hundred bp [56]. Though we have no direct assessment of the rate of outcrossing in *S. sclerotiorum*, the very small LD_2_ rate we observed suggests that it may be relatively frequent.

Like other outcrossing species, recombination was not uniform across the genome. In four non-structured subsamples comprising respectively 23, 13, 15 and 34 individuals (Figure 2 C), which we refer to as population-1, population-2, population-3 and population-4, we identified 384 recombination hotspots (Supplementary Table 4). Like other outcrossing species, we found that recombination rate was higher towards the ends of chromosomes where chromatin is more likely to be relaxed (Figure 2 D). Furthermore, recombination rate was negatively correlated with coding sequence density (Spearman’s ρ = -0.15, P = 0, Figure 2 E, Supplementary Figure 2) and recombination hotspots had a lower gene density than other regions (P < 0.0001, Figure 2 F). Though recombination rate was weakly negatively correlated with coding sequence density, the relationship between the two variables was not monotonic. Instead, there was an optimal gene density at which recombination rate peaked before declining rapidly (Figure 2 E, Supplementary Figure 2).

Our data suggest that recombination rate is generally higher outside of genic regions but low in the most gene-sparse parts of the genome. This is consistent with observations in other outcrossing species [57] where meiotic recombination within genes is selected against as it can lead to polymorphisms due to erroneous double strand break (DSB) repair, though meiosis is repressed in the most gene-sparse regions, which also tend to be heterochromatic.

Across most chromosomes and all four population samples, there were clear recombination coldspots that coincided with a single prominent drop in GC content and a single prominent spike in cytosine methylation based on bisulfite sequencing data generated in this study (Figure 2 G-H, Supplementary File 1, Supplementary Table 5). Decreased GC content and increased cytosine methylation are both hallmarks of eukaryotic centromeres [58,59], around which meiotic recombination is typically suppressed [60]. The convergence of these three observations, and previous predictions from optical mapping data [61], suggest that these sites are the centromeres of the *S. sclerotiorum* chromosomes, and, as in other outcrossing species, meiotic recombination is suppressed around them.

With the rapid decay of linkage disequilibrium, the presence of recombination hotspots, and the conspicuous recombination-related features characteristic of eukaryotic meiosis, we infer that *S. sclerotiorum* maintains genetic diversity across numerous populations through sexual outcrossing. While clonal lineages may endure over extended periods via self- fertilization, the ongoing process of sexual recombination among these lineages may be important for creating genotypic diversity. Presently, meiotic exchange is cryptic, as laboratory observations of sexual outcrossing are, to our knowledge, lacking.

Ecological theory suggests that loss of sexual reproduction initiates the gradual accumulation of deleterious alleles inseparable from beneficial ones, a phenomenon known as ’Muller’s ratchet’. Consequently, strictly clonal populations are rare, with most facing a trajectory toward extinction. Given the continuing pressure on *S. sclerotiorum* for survival across numerous host species, coupled with its apparent lack of host preference, it is not surprising that it exhibits many attributes indicative of sexual outcrossing. Drawing from our findings and those of others [37,43,44], we suggest that homothallism in *S. sclerotiorum* not only supports persistence of certain clonal lineages but also fosters universal sexual compatibility.

### The *Sclerotinia sclerotiorum* pan-genome graph suggests transposable elements create hotspots of structural diversity

To capture diversity of structural variation across the genome, we developed a statistic called ‘SVπ’. Akin to nucleotide [62] and synteny diversity [63], this statistic captures the average number of SVs per Kb between all pairs of individuals. We found that SVπ was positively skewed when calculated for 50 Kb sliding windows across the genome (Figure 3 A). This suggests that the *S. sclerotiorum* genome is mostly stable, with a few regions of excessive structural variation. We defined SV hotspots as sliding windows with a SVπ value above the 95^th^ percentile across the genome. Interestingly, more hotspots were detected on some chromosomes than others. For example, chromosome 12 contained six hotspots and had an average SVπ of 0.016, whereas, despite being larger, chromosome 6 contained only one hotspot and had an average SVπ of 0.009 (Figure 3 D, Supplementary Figure 3).

**Figure 3.**
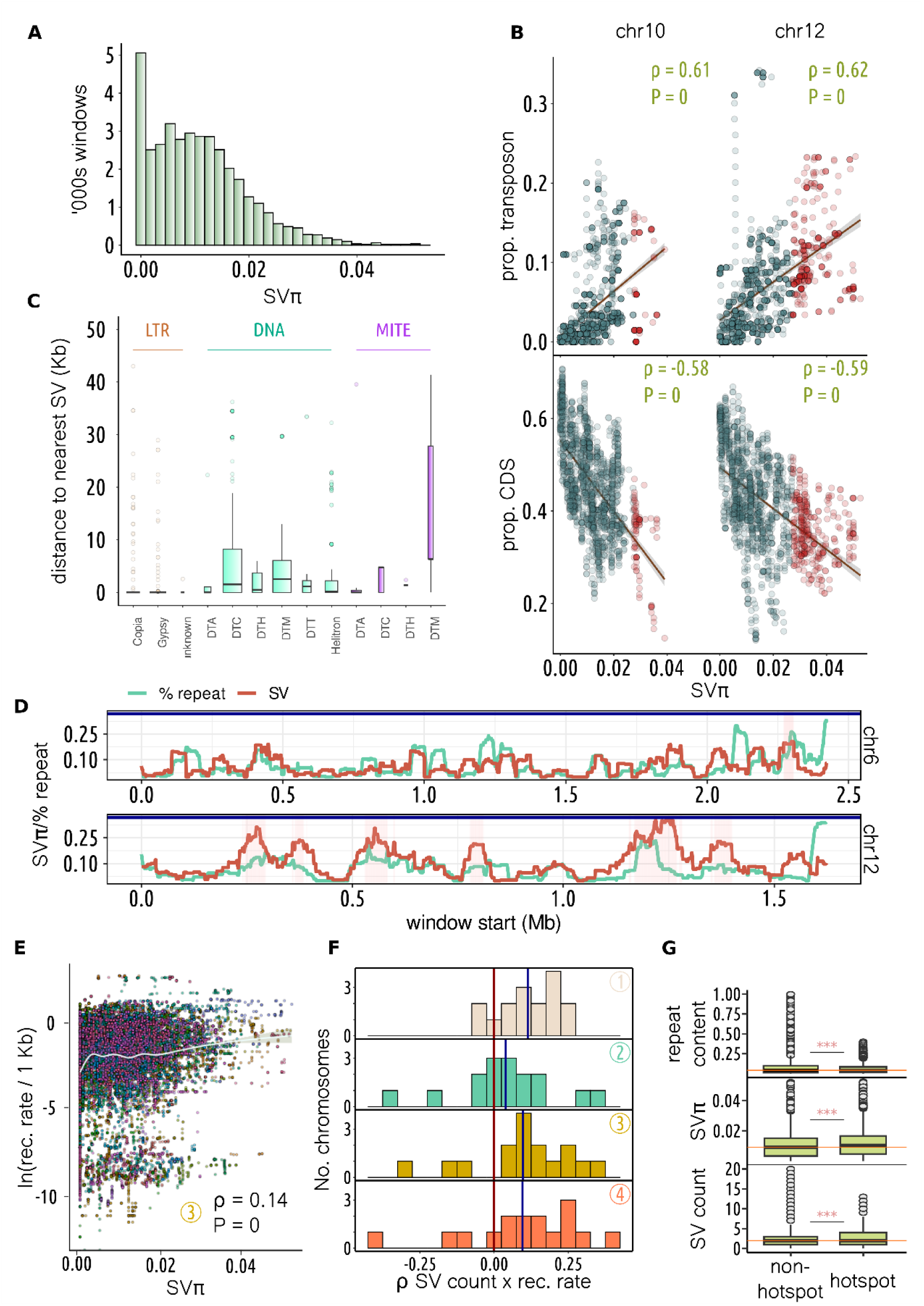
Analysis of structural variation across the *Sclerotinia sclerotiorum* pan-genome. **A** Distribution (y axis) of SVπ (x axis) for 50 Kb sliding windows. **B** For chromosomes 10 and 12, correlation between SVπ (x axis) and proportion transposon (top y axis) or coding DNA sequence (bottom y axis). Spearman’s ρ and P value depicted top-right. Blue lines show linear regression of y onto x and the shaded area 95 % confidence interval. Red points are SVπ hotspot (> 95^th^ percentile) windows. **C** The y axis shows distance to nearest structural variant (SV) for transposon families. Transposon classification is indicated at the top and family on the x axis. Boxes and whiskers show interquartile range. LTR retrotransposons were generally closer than other transposons to SVs (Kruskall-Wallis test show in Supplementary Table 6). **D** The y axis is SVπ or percent repeat for 50 Kb windows (scaled for visualisation). The x axis shows window start (Mb), and plots show chromosomes 6 and 12, the latter having the highest average SVπ and the most hotspots (shaded in pink). **E** Correlation between log recombination rate per Kb (y axis) and SVπ (x axis) across 50 Kb sliding windows. Chromosomes are plotted in different colours and data shown are for population-3. Spearman’s ρ was 0.14-0.15 for all populations (P = 0) but varied between chromosomes. **F** Distribution across chromosomes (y axis) of Spearman’s ρ for number of SVs and recombination rate in 50 Kb sliding windows. Though correlation strength varied between chromosomes, correlations were generally positive. **G** The y axis shows repeat content (top), SVπ (middle) and number of SVs (bottom) for windows that did not (left) and did (right) contain recombination hotspots. Boxes and whiskers show interquartile range; differences were significant according to a t-test (*** = P < 2.2e^-16^).

Transposable element and gene content were positively (Spearman’s ρ = 0.28, P = 0) and negatively (Spearman’s ρ = -0.33, P = 0) correlated with SVπ, respectively. The correlation between transposon/gene content and SVπ varied between chromosomes, with the highest correlations observed on chromosomes 10 and 12 (Spearman’s ρ = 0.61 and 0.62, Spearman’s ρ = -0.58 and -0.59, respectively (P = 0)) (Figure 3 B; Supplementary Table 8). The association between SVs and transposable elements was further supported by the observation that transposable elements were significantly closer than randomised loci to the nearest SV across all genomes (P = 0). Transposable elements in the long terminal repeat (LTR) family were significantly (P < 0.0001) closer to the nearest SV than those in eight out of 10 other families (Figure 3 C, Supplementary Table 6), suggesting they may strongly contribute to genome instability in *S. sclerotiorum*.

LTRs are a type of retrotransposon, which are transposons characterised by a copy and paste proliferation mechanism that involves transcription into RNA, reverse transcription into DNA and re-insertion into the genome [64]. Retrotransposons are unique to eukaryotes [65], and their replicative ability has made them often the dominant transposon class in eukaryote genomes [66]. Several studies have linked retrotransposons with virulence evolution in plant pathogens [67,68], including the host generalist species *Botrytis cinerea*, where they have been shown to encode small RNA effectors [69]. Our observations that retrotransposons are most strongly linked of all transposon classes to SVs suggests that they are the most active mobile elements in *S. sclerotiorum*. Their ongoing contribution to structural variation may be important for genomic plasticity in this species.

Stable, gene-dense and repeat-poor, and unstable, gene-sparse and repeat-rich genomic regions are common across eukaryote genomes [70]. The accumulation of transposons and SVs in gene-sparse regions is likely a result of relaxed selective pressure and accumulation of largely selectively neutral alleles. These regions can be important for adaptive evolution because they harbour extensive diversity in gene content and gene sequences [71]. When the environment changes, previously selectively neutral mutations may confer an advantage, leading to ongoing maintenance of these regions, and the transposons within them, in populations [72]. Our data show that, like those of most eukaryotes, the *S. sclerotiorum* genome is also partitioned into stable and unstable regions, and unstable regions are likely most strongly shaped by LTR retrotransposons. Overall, transposon content in the 25 *S. sclerotiorum* genomes was relatively low at 5.51 to 6.91 % (Supplementary Table 7). Despite this, transposable elements are responsible for creating considerable diversity in SVs across the *S. sclerotiorum* genome.

### Recombination and transposable elements make distinct contributions to structural variation

Several studies have shown that besides transposition, structural variation can be caused by recombination. However, little is known about the overall impact of recombination on structural variation in natural populations. In *S. sclerotiorum*, we found an overall correlation between SVπ and recombination rate for all four population samples we used for recombination rate estimation (Figure 3 E, Spearman’s ρ = 0.14-0.15, P = 0). Though this is suggestive of a link between SV diversity and recombination, it does not necessarily imply that recombination creates SVs, as this relationship could also be caused by increased haplotype diversity in regions with a high recombination rate. Therefore, to determine whether genomic regions with a high recombination rate may be more prone to development of SVs, we assessed the correlation between recombination rate and the overall number of SVs called against the reference genome. Though the strength of correlation between these parameters varied considerably between chromosomes and populations, we found that, on average, there was a weak to moderate correlation between total number of SVs and estimated recombination rate (Figure 3 F, mean Spearman’s ρ = 0.09). For 12, 8, 13, and 12 out of 16 chromosomes, for the four respective populations, there was a significant positive correlation between recombination rate and total number of SVs (P < 0.05, Supplementary Table 8). In contrast, only 1-3 chromosomes displayed a significant negative correlation between recombination rate and number of SVs.

Despite the correlations between recombination rate and both SVπ and total SVs across chromosomes and populations, there were far fewer instances of a positive correlation between recombination rate and transposon content, and the overall average of all Spearman’s ρs was close to zero at -0.0018 (Supplementary Table 8). Furthermore, recombination hotspots had a slightly but significantly lower average repeat content than other parts of the genome (5.71 % vs 6.97 %, P < 2.2e-16), despite having elevated SVπ (average of 0.012 vs 0.010, P < 2.2e-16) and more SVs (2.65 vs 2.40, P < 2.2e-16) (Figure 3 G). This suggests that meiotic recombination and transposition make orthogonal contributions to structural variation. In agreement, we found that the number of SVs was better described in a regression model by both average recombination rate and transposon content than transposon content alone, though transposon content was the dominant predictor in the model (likelihood ratio test P < 2.2e-16, transposon *F* = 74.86, recombination rate *F* = 13.08).

Our analyses document an interesting link between estimated recombination rate and the rate of structural variation in the *S. sclerotiorum* genome. This is not surprising given the mutagenic properties of meiosis. Given the relatively low level of transposable element content in the *S. sclerotiorum* genome, recombination through meiotic exchange could be an additional important source of structural variation. Our regression model suggests that recombination rate is far outweighed by transposon density as a predictor of genome stability. However, since recombination rate was typically higher in regions of intermediate gene density, recombination may have a greater chance of inducing SVs that impact gene function.

### *Sclerotinia sclerotiorum* has a closed pan-genome with relatively few non-syntenic blocks of genes

The gene-space within a pan-genome lies on a spectrum from high variability in certain species to remarkable stability in others. Species harbouring a limited number of dispensable genes are characterised by closed pan-genomes, while those with diverse gene content are classified as having open pan-genomes [73]. To assess the openness of the *S. sclerotiorum* pan-genome, we sampled from two to all 25 strains in our dataset and plotted number of strains against number of novel genes. We found that the number of additional genes brought by adding a new strain plateaued quickly at 5-10 strains, indicating that most dispensable genes in the population are present in multiple strains (Figure 4 A).

**Figure 4.**
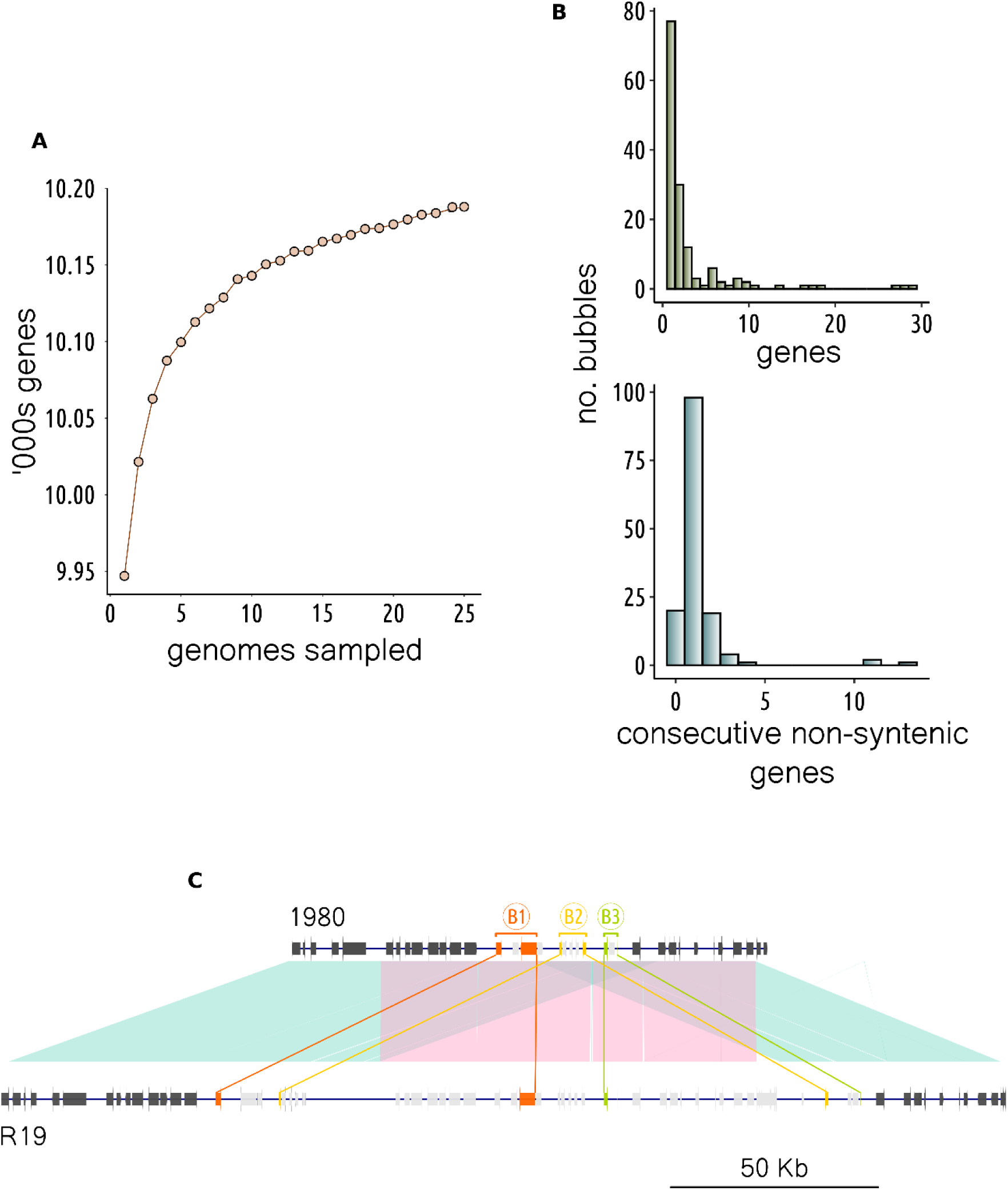
Gene content variability in the *Sclerotinia sclerotiorum* pan-genome. **A** The relationship between total number of unique genes (y axis) and number of genomes sampled (x axis). **B** Number of gene bubbles (y axis) and number of genes they contained (top) or number of consecutive missing genes they contained (bottom). **C** A region in the 1980 reference genome that had a complex rearrangement in the isolate R19 and no other isolates. This region contained the largest three gene bubbles, indicated here with B1 (orange), B2 (yellow) and B3 (green). Start and end genes for each called bubble are indicated in their respective colours and non-syntenic genes within bubbles are in light grey. Neighbouring genes are in dark grey. The shaded area connects homologous regions and the pink region is duplicated in R19.

Across all strains, we identified 10,188 unique genes, of which 553 (5.43 %) were dispensable. Few informative Gene Ontology terms were over-represented among these genes, although we noted an over-representation of the term ‘GO:0031177’ (P = 0.012), which is ascribed to genes containing a phosphopantetheine binding site. Since one of the main functions of this site is in secondary metabolite biosynthesis by non-ribosomal peptide synthases (NRPSs) and polyketide synthases (PKSs), it is not surprising that we also found an over-representation of genes in secondary metabolite biosynthesis clusters among dispensable genes (odds ratio = 2.28, P = 1.44e-11). We found no significant over-representation of secreted proteins, regardless of size (odds ratio = 0.73, P = 0.18 for >= 300 amino acids; odds ratio = 1.38, P = 0.19 for <= 300 amino acids).

Based on a graphical representation of gene synteny, we identified 615 runs of one or more genes that were non-syntenic between strains (Figure 4 B). In keeping with graph terminology, we refer to these as ‘gene bubbles’. The number of genes in gene bubbles ranged from 1 to 34, with most gene bubbles containing only a single gene (Figure 4 B). Consecutive runs of missing genes within bubbles ranged from 0 (i.e. the bubble was an inversion) to 13 (median = 1) (Figure 4 B). The largest three consecutive runs of missing genes within bubbles were identified on chromosome 12, which was the chromosome with the highest SVπ. Closer inspection of these runs identified a complex region partially duplicated in the strain R19, which was sampled in 2007 from buttercup in Warwickshire in the UK (Figure 4 C). Many of the genes in this region were likely transposon genes, as they were annotated with Pfam domains such as RNase H (PF00075), reverse transcriptase (PF00078), and endonuclease (PF14529) (Supplementary Table 9). However, there were also glycosyl hydrolases (PF00722), ubiquitins (PF00240) and a major facilitator superfamily transporter (PF07690). The fact that this region was the same in all strains apart from R19 could mean it is deleterious. Alternatively, it could be a relatively new polymorphism whose evolutionary fate has not yet been determined. So far, the polymorphism does not appear to be detrimental to infection on brassicas, since R19 is more aggressive than several other diverse isolates from the UK [74].

The closed *S. sclerotiorum* pan-genome contrasts the pan-genomes of host specialist fungal pathogens. For instance, in a population sample of 19 global isolates of *Z. tritici*, approximately 40 % of genes were dispensable [8], and in 26 strains of the wheat pathogen *Pyrenophora tritici-repentis* 43 % [75].

To our knowledge, little is known about what shapes pangenome openness in eukaryotes. However, ecological theory suggests that selective pressure from the host is stronger on host specialists than generalists [33]. To our knowledge, there are no *S. sclerotiorum* strains unable to reproduce on a single host species or genotype. It is unlikely, therefore, that a single virulence gene, such as an effector, would ever confer a strong host-driven selective advantage in this species. Therefore, maintenance of a repertoire of dispensable virulence proteins to ensure adaptability to a constantly changing host environment seems unlikely. Instead, the closed pan-genome of *S. sclerotiorum* aligns with previous research suggesting that it, and other host generalists, have evolved toward energetic optimisation of core virulence genes that function on multiple host species [31,34].

### Structural variation may have a strong impact on adaptive flexibility of life history traits

Adaptive flexibility and fitness of a population are underpinned by genotypic variation that impacts life history traits. As a global host generalist agricultural pest, *S. sclerotiorum* is exposed to diverse environments and must be adaptable to a range of temperatures and stressors, such as host metabolites. To assess global phenotypic diversity in *S. sclerotiorum*, we measured 14 life history traits across 103 genotypically distinct strains, including relative growth on the Brassicaceae defence compounds brassinin and camalexin, the Fabaceae defence compound medicarpin, the reactive oxygen species H_2_O_2_ (ROS), and the two fungicides azoxystrobin and tebuconazole; growth and relative growth at 15, 20 and 25 °C; and fecundity-related traits including number, and average and total weight of sclerotia.

We found significant differences between isolates from different geographical origins for eight of these traits. Both European and Australian strains grew faster at 15 °C than Canadian strains (Figure 5 A) (P = 0.014 and 0.007, respectively). At 20 °C, European strains grew significantly faster than Australian but not Canadian strains (P = 0.003 and 0.52, respectively), though Canadian strains grew at a similar rate to Australian strains (P = 0.40). At 25 °C, European strains grew faster than both Canadian and Australian strains (P = 0.035 and 0.00072, respectively). Relative growth (growth divided by growth at 20 °C, generally considered the middle of the optimum range [76,77]) at 15 °C was significantly lower for both European and Canadian strains compared with Australian strains (P = 0.035 and 0.049, respectively), though relative growth at 25 °C was not significantly different between strains from different global regions.

**Figure 5.**
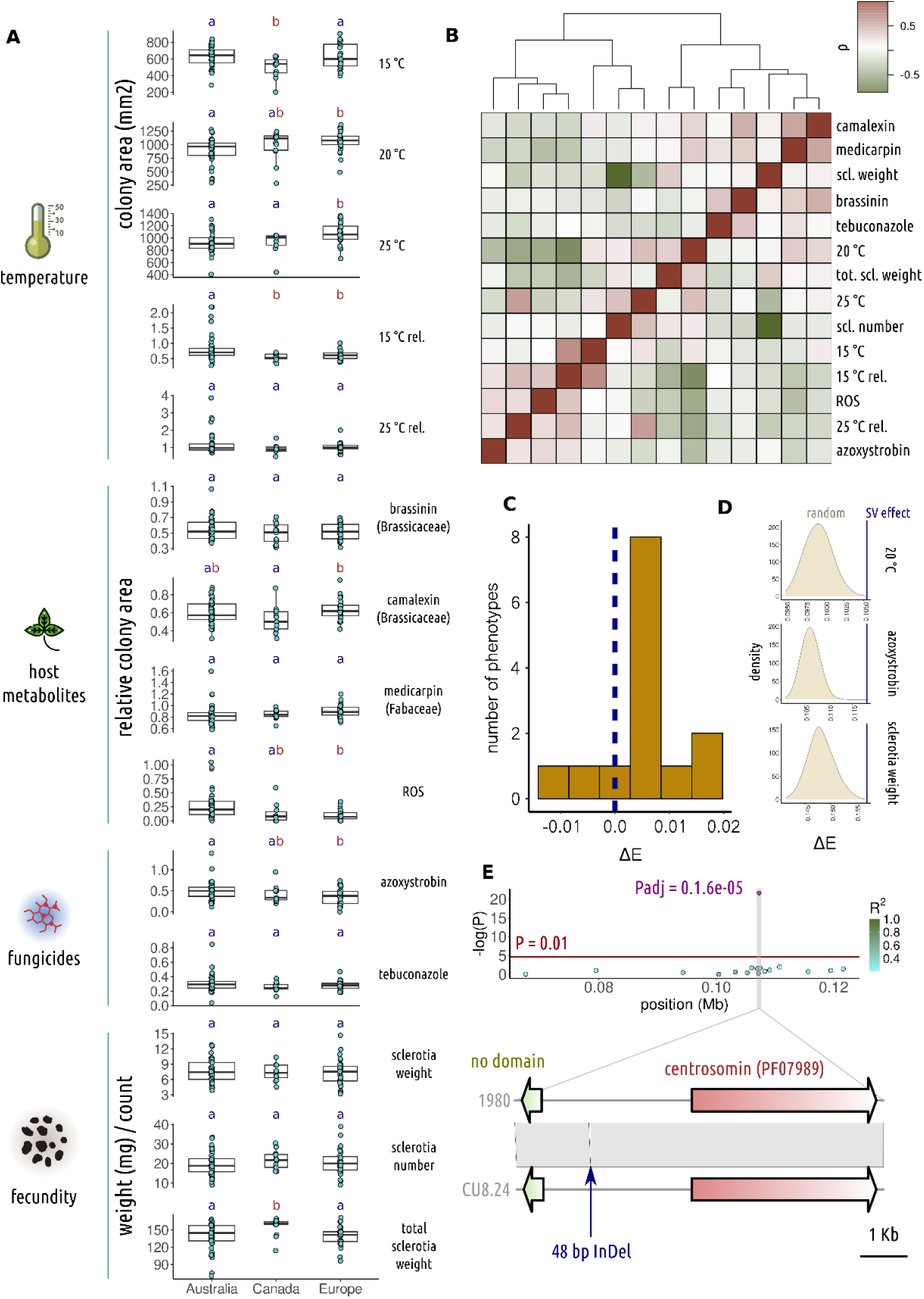
Life history traits assessed across a subset of *Sclerotinia sclerotiorum* strains. **A** Boxplots of measurements for life history traits (indicated to the right of the plot) in four categories (to the left). The distribution is plot for strains from the three major geographical regions, Australia, Canada and Europe. Points are the individual data points and box and whisker plots show interquartile range. The letters a and b above plots indicate significant differences between groups. **B** The top panel is a heatmap (rows are in the same order as columns), showing Pearson’s ρ between measurements for the 14 life history traits. Colouring goes from green (negative correlation) to red (positive). The dendrogram shows hierarchical clustering of the traits. **C** The distribution (y axis) across the 14 traits of mean effect size of non-SVs subtracted from mean effect size of SVs (ΔE), where effect sizes are absolute. **D** The y axis shows the density of measurements of absolute effect size for 500 random samples with an identical minor allele frequency distribution to that of SVs. The blue line shows the observed absolute effect size for SVs. For these three traits, the P value of this test was 0. **E** A region surrounding a quantitative trait locus (QTL) for relative rate of growth on azoxystrobin (top), with -log(P) on the y axis and position in Mb on the x. The colours of points represents linkage disequilibrium of variants at the different positions with the QTL (the purple point with associated P value). The red line is a P value of 0.01. Below this region, the two genes neighbouring the 48 bp InDel underlying the QTL are illustrated. These included a small gene with no known domains and a larger centrosomin-encoding gene.

Differences in growth rate at different temperatures between these populations could be a result of adaptation to prevailing temperatures during the growing season for major host crops, a phenomenon that has been previously observed at a local level in Australia [50,51,78]. However, it is difficult to completely align our observations with the likely reproductive cycle of *S. sclerotiorum* in these three global cropping regions, as different host crops are likely to be available at different times of year. For example, the major host species *Brassica napus* usually flowers in spring in the UK, where temperatures are often lower than when *B. napus* flowers in Western Australia in July. On the other hand, some hosts, such as lettuce, may be also present later in the season in the UK. A weaker adaptation to lower temperatures is possible for Canadian strains, which would likely infect *B. napus* when it is flowering during the hotter summer months.

Among host antimicrobial metabolites, we found a significant increase in growth in European strains compared with Canadian on camalexin (P = 0.015) and a significant decrease on ROS (0.0015) compared with Australian strains. Growth on ROS was lower for both European and Canadian compared with Australian strains, though the range in growth of Canadian strains meant the difference between Australian and Canadian strains was not significant (P = 0.12). Similarly, European strains had a significantly lower growth rate on azoxystrobin compared with Australian strains (P = 0.035), and Canadian strains were in the middle (P = 0.69 and 0.64 compared with Australian and European strains, respectively).

Interestingly, Canadian strains produced a greater total sclerotia weight compared with strains from Australia and Europe (P = 0.034 and 0.036, respectively). This seemed to be due to a slight increase in the mean of both sclerotia number and weight. The size of sclerotia has been previously linked with the rate of germination [79] and number of apothecia per sclerotium [80], suggesting it is an important component of fecundity. Phenotypic variation in this trait may have important implications for pathogen proliferation and epidemic potential of different populations.

The traits we measured had complex genetic synergisms and antagonisms with one another, for instance brassinin and camalexin tolerance were positively correlated (Pearson’s ρ = 0.43) and shared positive genetic covariance (0.93) (Figure 5 B, Supplementary Table 10). The same was true of camalexin and medicarpin tolerance (Pearson’s ρ = 0.48, genetic covariance = 0.65). Other traits were negatively correlated and had negative genetic covariance, for instance growth at 20 °C and azoxystrobin tolerance (Pearson’s ρ = -0.33, genetic covariance = -0.97), and total sclerotia weight and relative growth at 15 °C (Pearson’s ρ = -0.39, genetic covariance = -0.63). This suggests that complex trade-offs and synergisms between life history traits may influence fitness [5].

Several studies have shown that SVs have a major role in creating phenotypic diversity, and graph pan-genomes in which SVs can be reliably called have shown that they can be a major component of missing heritability [19,81]. To test the relative impact of SVs on life history traits, we conducted a genome-wide association study (GWAS). For all traits, quantile-quantile plots suggested that the model we used adequately controlled P value inflation due to population structure (Supplementary Figure 4).

We found that the average absolute effect size of SVs was higher than that of non-SVs across 11 of the 14 traits, significantly higher for eight (P < 0.05), and lower for 2 of the 14 traits, tebuconazole and ROS tolerance. Notably absolute effect size was on average 0.015 and 0.020 points higher for relative growth on azoxystrobin and total sclerotia weight, respectively (P < 0.0001, Figure 5 C, Supplementary Table 11). On average, SVs had a lower average minor allele frequency than other variants (0.18 vs 0.21), which could lead to an increase in the variance of effect size estimates. Therefore, we took 500 random samples of non-SVs of equivalent size and minor allele frequency distribution to SVs and assessed how many times their average absolute effect size was more than or equal to that of the SVs. Based on this test, azoxystrobin tolerance, growth rate at 25 °C and average sclerotia weight, were impacted more strongly by SVs than other variants in 100 % of random samples (P = 0), as well as having significant increases for the first test (P < 0.005) (Figure 5 C, Supplementary Table 11). Three other traits, medicarpin tolerance, growth at 25 °C and total sclerotia weight were more strongly impacted compared with more than 90 % of random samples (P < 0.1), as well as showing a significant increase according to the first test (P < 0.005). According to the randomisation test, ROS and tebuconazole tolerance were also significantly more weakly impacted on average by SVs than other variants (P = 1 – all randomisations had a higher mean absolute effect size). We infer from this analysis that SVs could have a larger impact on many of these traits than other variants, though genetic architecture with respect to SVs likely varies considerably between traits.

As an alternative to assessing the absolute effect size, we performed regressions on different genomic relationship matrices, which either included or did not include all variants in linkage disequilibrium with SVs. Using cross-validation, we found that for medicarpin and ROS tolerance, growth at 15 °C and 20 °C, and sclerotia number, models that included variants in LD with SVs had a better predictive ability than models that did not (Pearson’s ρ = 0.25 vs 0.23, 0.38 vs 0.37, 0.03 vs 0.02, 0.34 vs 0.29 and 0.14 vs -0.14, respectively (Supplementary Table 12)). Given absolute effect size for ROS tolerance was lower on average among SVs, it is possible that although individual SVs have a relatively weak impact on this trait, as a collective they explain a relatively larger proportion of additive heritability. For the other traits, the improvement in predictive ability was in accordance with the increase in absolute effect size according to at least one of the analyses mentioned previously.

Overall, we identified 15 variants with a significant (Benjamini-Hochberg adjusted P < 0.05) impact on phenotype across six traits. 13 of these variants were intergenic SNPs or InDels, one was a synonymous SNP and the other was a disruptive in-frame InDel. The genes associated with these variants had diverse functions, which may be speculatively associated with each of the traits (Supplementary Table 13). Though none of these variants were SVs or in linkage disequilibrium with neighbouring SVs, one of them was a 48 bp InDel, close to the 50 bp cutoff we used for designating variants as ‘structural’. This variant, at position 107,298 on chromosome 12, was a deletion that significantly increased relative growth on azoxystrobin (P = 1.6e^-5^) (Figure 5 E). There were two genes either side of this variant on opposite strands, one encoding a 78 amino acid protein with no known domains and the other a 1,516 amino acid protein containing a centrosomin domain (PF07989). The variant was, respectively, 1,177 and 2,571 bases away from the transcription start sites of the shorter and longer genes. The amino acid sequences of both proteins were well-conserved in fungi, with 80-100 % similarity to homologues from species in the Helotiales, suggesting they are genuine genes (Supplementary Table 13). With the current data, it is not possible to determine which, if either, of these genes’ functions is linked with growth rate on azoxystrobin. However, in fungi, centrosomins are localised to spindle pole bodies, which are structures analogous to animal centrosomes, the main sites of coordination of microtubule activity during mitosis [82]. It is conceivable that the relative rate of cell division on a particular stressor could be impacted by mutations affecting genes encoding the cell division machinery.

Our analyses are in accordance with several studies showing that SVs may have an outsized impact on phenotype [19,83,84]. In our analyses, which used a GWAS approach, we were restricted to common (minor allele frequency > 0.05), biallelic SVs. Therefore, we likely favoured less deleterious SVs, since deleterious SVs with the strongest phenotypic impacts are usually rare in populations [84,85]. Despite this, we observed an overall stronger impact of SVs than other types of variants on many life history traits. This was also despite the observation that SV diversity tended to cluster in polymorphic, repeat-rich genome regions, which can often be sites of selective neutrality. Since the *S. sclerotiorum* genome is relatively gene-dense, containing few repeat-rich regions, this could suggest that the amount of selectively neutral structural variation it contains is relatively low. Alternatively, the variation across many of the traits we assessed could be selectively neutral or perhaps deleterious. In this case, the stronger link between SVs and phenotype is indicative of SVs underlying a particularly large amount of adaptive potential.

## Conclusion

Collectively, our results portray *S. sclerotiorum* as both a clonal and sexually outcrossing pathogen with limited diversity in gene content. Despite this limited diversity, *S. sclerotiorum* isolates vary considerably in life history traits. SVs may make a particularly strong contribution to this variability for some traits, and are likely generated through two distinct mechanisms, meiotic recombination and transposition, the latter being the dominant mechanism.

The limited genic diversity of *S. sclerotiorum* contrasts the highly variable open pan-genomes of many host specialist species. Such stable gene content aligns with the hypothesis that *S. sclerotiorum*, a niche generalist, is a ‘jack of all trades’, with a core, multifunctional infective arsenal enabling fitness on hundreds of host species.

At this stage, the relative importance of meiosis and transposition in the generation of adaptively advantageous SVs is unknown. Given the likely general evolution of *S. sclerotiorum* towards a stable, repeat-poor genome, and the stronger contribution of SVs to variability in life history traits, it is possible that meiosis, through its tendency to create SVs, may have a significant role in adaptation beyond its well-recognised role in recombination of alleles into new haplotypes.

Overall, our data shed considerable light on the evolutionary processes at play in an important host generalist plant pathogen of agricultural significance.

## Methods

### Assessment of life history traits

A sclerotium of each isolate was cut in half, placed on a potato dextrose agar (PDA) plate, and incubated at 20 °C for 4 to 5 days. Hyphae from the leading edge of the mycelium were cut with a 3-mm cork-borer, placed onto fresh PDA plates and incubated at 20 °C for 2 days to source actively growing mycelium. Actively growing mycelia were subcultured with a 3-mm cork-borer onto appropriate PDA plates for trait assessments.

To measure the effect of temperature on mycelium growth, mycelia were subcultured onto PDA and grown at 15 °C, 20 °C and 25 °C for 1 day. To measure the effect of host metabolites on mycelium growth, mycelia were subcultured onto PDA supplemented with 50 µM brassinin (Sigma-Aldrich), 20 µM camalexin (Sigma-Aldrich), 20 µg/mL medicarpin (TargetMol), or 200 µg/mL hydrogen peroxide H_2_O_2_ (Westlab) and grown at 20 °C for 1 day. Brassinin, camalexin and medicarpin were all dissolved in DMSO prior to PDA supplementation. To measure the effect of fungicides on mycelium growth, mycelia were subcultured onto PDA supplemented with 0.2 µg/mL azoxystrobin and 50 µM salicylhydroxamic acid (SHAM), or 0.16 µg/mL tebuconazole and grown at 20 °C for 1 day. Azoxystrobin and tebuconazole were dissolved in ethanol and SHAM was dissolved in water prior to PDA supplementation. All *S. sclerotiorum* strains were also grown on PDA supplemented with the equivalent concentration of DMSO, ethanol, or ethanol and SHAM as used for the aforementioned compounds. Though tebuconazole was dissolved in ethanol, ethanol control plates were not available for the experiment, so growth on tebuconazole was normalised to growth on DMSO. We did this because neither DMSO nor ethanol strongly impacted growth, whereas growth rates were often variable between experiments. For mycelium growth measurements on PDA, photographs were taken of each inoculated PDA plate. The colony area was measured using Image J software. Colony area relative to growth at 20 °C on PDA (plus appropriate solvent or SHAM) was calculated for each isolate.

To measure the effect of temperature on sclerotia formation, mycelia were subcultured onto PDA and grown at 20 °C for 1 month. Mature sclerotia were then air-dried for 3 days. The number and weight of sclerotia per plate were recorded.

### DNA extraction and sequencing

To extract high molecular weight DNA, sclerotia were cut with a sterile scalpel and placed with the cut side touching the surface of the medium on potato dextrose agar (PDA) plates. After three to four days at room temperature in darkness, strains were sub-cultured onto fresh PDA plates from agar plugs using a sterile cork borer and forceps. After two further days at room temperature in darkness, strains were sub-cultured again by placing four plugs for each strain into 100 ml of potato dextrose broth (PDB) in 250 ml conical flasks. These liquid cultures were incubated at room temperature with ambient light conditions in the laboratory with shaking at 150 rotations per minute (RPM) for three days and used to generate protoplasts.

Protoplasts were generated by removing fungal cultures that had grown around plugs and placing two plugs each in 250 ml conical flasks with 40 ml enzymatic digestion solution containing 0.8 M mannitol, 200 mM citric acid/tri-sodium citrate buffer and 1.5 % w/v lysing enzymes from *Trichoderma harzianum* (L1412, Sigma, now discontinued). Digestions were incubated for three hours at 30 °C with shaking at 80 RPM. All protoplasts from each conical flask were then filtered through a 100 µm cell strainer (CLS431752, Merck) into one 50 ml falcon tube and pelleted using a swinging bucket rotor centrifuge at 1,000-2,000 x g for 2-3 minutes at 4 °C. Protoplast pellets were re-suspended in 200 µl Tris-EDTA (pH 8.0).

The resuspended protoplasts were then used as input for the MagAttract high molecular weight DNA extraction kit (67563, Qiagen), which was used with the manufacturer’s protocol for blood cells with the following modifications: 80 µl proteinase K was used instead of 20 µl, 20 µl of RNase A was used instead of 4 µl, 600 µl of buffer AL was used instead of 150 µl, 25 µl of MagAttract Suspension G was used instead of 15 µl, 600 µl of buffer MB was used instead of 280 µl; before adding MagAttract Suspension G, samples were also filtered through miracloth to remove debris. High molecular weight DNA was then sequenced on an Oxford Nanopore MinION using an SQK-LSK109 library prep kit multiplexed with the native barcoding expansion pack EXP-NBD104 on a R9.4.1 version flowcell.

To extract DNA for Illumina sequencing, the same procedure was used for initial culturing of *S. sclerotiorum* strains. Cultures from PDB were then snap frozen in liquid nitrogen and freeze-dried overnight. Portions of approximately 1 g of freeze-dried samples were then cut with a sterile scalpel and placed using forceps into 2 ml screw-capped Eppendorf tubes with a single ball bearing. To each tube, 700 µl lysis buffer (50 mM Tris-HCL, 50 mM EDTA, 3 % sodium dodecyl sulfate, 1 % 2-mercaptoethanol) was added, and samples were ground in a MiniG model 1600 at 15000 RPM for 2 minutes. Samples were then centrifuged at 17000 RPM for one minute, and ball bearings were removed. To each tube, 100 µl RNase A was added and tubes were then incubated for 1 hour at 65 °C. To each tube, 700 µl chloroform:phenol (50:50) was added and tubes were vortexted. Tubes were then centrifuged at maximum speed for 5 minutes before removal of the aqueous phase. Then, 700 µl of chloroform:isamyl alcohol was added, the tubes vortexed and centrifuged again at full speed for 5 minutes. The aqueous phase was again removed and DNA was then precipitated using 6 M sodium acetate. Paired end Illumina sequencing was conducted at Genomics WA on a NovaSeq 6000 flowcell at 2 x 150 cycles to yield 1.2 Gb per sample. For genomic DNA methylation analysis, *S. sclerotiorum* 1980 (ATCC 18683) was propagated on minimal salts – glucose (1% w/v) (MS–Glu) agar. The inoculum for all experiments was prepared by grinding 2 g of sclerotia in 200 mL of MS–Glu in a Waring blender for 4 min. The volume was increased to 500 mL in a 1 L baffled flask and the culture incubated at 20 °C with shaking (60 r/min) for 3 days. 1 g of mycelia (wet mass) was spread over a 5-cm-diameter area of *B. napus* leaf surface and incubated in a humidified chamber. Leaves from 45-day-old plants were used. 3 biological replicates (3 different flasks of culture inoculated onto different leaves) were collected. The mycelial mat was collected from the lesion using forceps at 48 hours post-inoculation, plant material was removed, and the samples frozen immediately in liquid nitrogen. Samples were ground in liquid nitrogen using a mortar and pestle, then genomic DNA was extracted from a 100 mg sample using the DNeasy Plant Mini kit (Qiagen). Genome Quebec performed whole genome bisulfite sequencing using the NEB Next kit, then sequenced 2x250 bp on an illumina NovaSeq6000.

### Trimming and demultiplexing reads

FAST5 files from the Oxford Nanopore were basecalled using Dorado version 0.3.2 and de-multiplexed using Guppy version 6.5.7. Illumina whole genome sequencing reads were trimmed using cutadapt version 2.8 [86] with appropriate adaptor sequences. Bisulfite sequencing reads were assessed for quality and low quality bases and adapters were trimmed using CLC genomics workbench 20.0.2.

### Analysis of bisulfite sequencing data

Methylation analysis was performed using the CLC genomics workbench 20.0.2. Reads were mapped to the *S. sclerotiorum* genome (GCF_000146945.2) using “Map Bisulfite reads” (directional mapping and default mapping options). Methylated residues were identified using “Call Methylation Levels” with default settings with the following exceptions: exhaustive context-independent calls, minimum read depth of 10 reads. The data presented in the results section is a count of methylated bases per 50 Kb sliding window.

### Genome assembly

Genomes were assembled from Nanopore reads using Flye version 2.8.1-b1676 [87] and polished using Illumina reads, either from [29] or generated in this study, with one round of Polypolish version 0.5.0 [88], followed by one round of Pilon version 1.24 [89]. Before subsequent analyses, mitochondrial contigs were removed from assemblies using the following procedure. Within Geneious Prime version 21.2.2, the Minimap2 version 2.24 [90] plug-in was used to align a published *S. sclerotiorum* mitochondrial genome (NCBI accession KX351425) to each of the polished genomes. Contigs that aligned to this accession with more than 95 % identity were separated from nuclear contigs, which we focus on in this study.

Polished nuclear chromosomes for each genome were scaffolded to the *S. sclerotiorum* reference genome [30] using the command ‘scaffold’, with the flags ‘-u -w -o’, from RagTag version 2.1.0 [91]. We then used the following process to finalise the scaffolded assemblies. First, Nanopore reads for each assembly were self-corrected using Canu version 2.2 [92]. Within Geneious Prime version 21.2.2, corrected reads were then aligned to their respective assemblies using the Minimap2 version 2.24 plug-in and used to manually add telomeres and subtelomeric sequences to the ends of chromosomes where they could be recovered from reads. Gaps between scaffolded contigs were also removed if there was extensive read support for joining the contigs.

Commands from Mummerplot version 3.1 [93] were then used to check for misassemblies. First, ‘nucmer’ was used to align each assembly individually to the *S. sclerotiorum* reference genome with the option ‘--mum’. The ‘delta-filter’ command was then used, with the options ‘-1 -i 95 -l 10000 -u 100’ to filter the output of nucmer. The filtered output was then passed to the command ‘show-coords’ to produce coordinates of scaffold mappings to the *S. sclerotiorum* reference genome. The command ‘awk ’NR > 5 {strand="+"; if($2 < $1){strand="-"};print $12"\t"$1"\t"$2"\t"$13"\t.\t"strand}’’ was then used to convert these coordinates into browser extensible data (BED) format. In Geneious, the BED file containing alignments to 1980 and mappings of self-corrected reads were used to judge whether chromosome segments had been artificially joined by the assembler. For instance, if one chromosome was unusually large and contained two different segments mapped to different 1980 chromosomes, it was split in two if (i) very few reads supported the join and (ii) reads showed evidence of extensive soft clipping either side of the join. Where chromosomes were split, genomes were scaffolded a second time using RagTag and gaps between joined chromosome segments were removed if aligned reads supported the join. The majority of chromosomes had zero gaps after the first round of scaffolding and only two in each of three strains were broken and re-scaffolded based on the latter procedure.

### Genome annotation

For comparative purposes, all genomes, including the reference genome, were annotated with the same procedure. First, repetitive sequences were annotated using EDTA version 2.2 [94] with the flag ‘--anno 1’. Then, Braker3 [95] was used to annotate genes with both RNA sequencing and amino acid sequences as evidence with the additional flags ‘--fungus’, ‘--prot_seq=Fungi.fa’, ‘--august_args=”--species=botrytis_cinerea”’. The RNA sequencing data used for annotation were derived from 32 samples from the sequence read archive (SRA) detailed in Supplementary Table 14. Reads from these samples that were derived from infected plant tissue were first filtered by alignment to their respective host genomes (Supplementary Table 14) with Hisat2 version 2.1.0 [96] and keeping unmapped reads with ‘--un-conc’ for paired end reads or ‘--un’ for single end reads. Filtered reads, and reads not from plants, were then aligned to each of the *S. sclerotiorum* genomes with Hisat2, converted to bam format with samtools version 1.10 [97] ‘view’ and used as input for Braker3. Amino acid sequences from Braker3 annotations were combined into a non-redundant set of genes for all isolates using cd-hit version 4.8.1 [98], and non-redundant proteins were annotated with InterProScan version 5.54-87.0 [99]. Secondary metabolite clusters in this set of proteins were identified using antiSMASH version 7.0 [100], and secreted proteins were identified with SignalP version 6.0 [101].

### Pan-genome graph construction and variant calling

Using the 24 Nanopore assemblies and the *S. sclerotiorum* reference genome (GenBank reference GCA_001857865.1), a pan-genome graph genome was constructed with cactus 2.5.2 [102]. Illumina reads from [29] and those generated in the current study were mapped to the pan-genome GBZ formatted graph using the ‘giraffe’ command of vg version 1.52.0 [18]. The resulting GAM formatted files, one for each set of Illumina reads, were filtered using the vg command ‘filter’, with the flags ‘--min-primary 0.90 --frac-score --substitutions - -min-end-matches 1 --min-mapq 15 --defray-ends 999’. Filtered GAM files were then passed to the vg command ‘pack’ to create pack formatted read support files for each variant, with the flag ‘--min-mapq 5’. The ‘call’ command from vg was used to call variants from the pan-genome graph using the Illumina reads with the flags ‘--ploidy 1 --genotype-snarls’ and create a variant call format (VCF) file. The VCF files for all samples were combined into a single file by first converting them to gzip format with ‘bgzip’, then merging them with the bcftools version 1.10.1 [103] command ‘merge’, with the option ‘--all’. We then filtered this VCF with vcftools version 0.1.16 with the options ‘--minQ 30’ and ‘--minDP 5’. To do this, we had to first set all variants to ‘PASS’ because bcftools merge adds a filter to the whole variant if only a single sample is filtered in one of the inputs. We did this using a simple Awk script. After calling variants present in the pan-genome graph, additional variants present in Illumina reads but not in the 25 genomes that made up this graph were called using the following procedure. First, filtered GAM files were converted to binary alignment map (BAM) files using the vg command ‘surject’ and sorted using the samtools command ‘sort’. Then, the command ‘mpileup’ from bcftools was used with the flags ‘--max-depth 1000 --output- type u’ and the BAM files as input. The output of ‘mpileup’ was piped to the bcftools command ‘call’, which was run with the flags ‘--output-type v --multiallelic-caller --ploidy 1’ to create a VCF file. We then filtered this VCF using vcftools with the options ‘--minQ 30’, ‘-- minGQ 30’ and ‘--minDP 5’.

Finally, we used vcftools to remove variants called by vg from the VCF created using bcftools with the options ‘--min-alleles 2’, ‘--mac 1’ and ‘--exclude-positions’, and concatenated the resulting VCF with the one produced using vg with the bcftools command ‘concat’ with the option ‘--allow-overlaps’. We further filtered the final VCF with a Python script (Supplementary File 2) to remove variants with a missing call rate of >= 0.2.

### Population structure characterisation

To identify clones, a VCF containing variants called against the graph pan-genome was used with plink version 1.9 [104] to generate an identical by state relationship matrix, with the flags ‘--snps-only’, ‘--biallelic-only’, ‘--double-id’, ‘--geno 0.2’, ‘--mind 0.2’ and ‘--make-rel square 1-ibs’. Then, the matrix was used to construct a distance matrix and dendrogram using hierarchical clustering. Clones were identified based on a relatedness of 98 % identical by state with the R base function ‘cutree’.

Population structure was analysed using ADMIXTURE version 1.3 [105]. As for the identical by state relationship matrix, we considered only biallelic SNPs. These were first filtered using plink with the flag ‘--indep-pairwise 50 10 0.1’, and admixture was run for 1 to 10 ancestral populations with cross-validation. A scree plot was used to determine the most appropriate number of populations to use based on cross-validation error. Principal component analysis was also performed with plink using the flag ‘--pca 4’.

### Assessment of structural variant diversity

To assess the diversity of SVs across the genome, we developed a novel statistic that we refer to as 𝑆𝑉_𝜋_, which is calculated as follows. First, we calculate 𝑆𝑉_𝑛_, which is the sum of the number of SVs between all pairs of individuals, excluding self-comparisons.

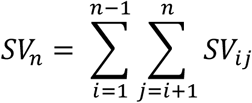

Where 𝑆𝑉_𝑖𝑗_ is the number of variants that are >= 50 bp in at least one individual for individuals 𝑖 and 𝑗 in the set of 𝑛 individuals in the sample. 𝑆𝑉_𝑛_ is then normalised in the following way to obtain 𝑆𝑉_𝜋_:

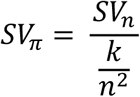

This divides 𝑆𝑉_𝑛_ by the number of possible pairs of individuals and the length of the sequence under consideration, 𝑘. Since 𝑘 varies between individuals depending on the SV alleles they contain, it is calculated in the following way:

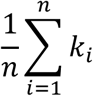

That is, 𝑘 is the average value of all 𝑘_𝑖_ sequence lengths, in Kb, in the set of 𝑛 individuals. The statistic is trivial for regions containing only biallelic SVs but more computationally challenging for regions with multi-allelic variants.

The statistic 𝑆𝑉_𝜋_ is an estimate of the average number of SVs that are present per Kb between all pairs of individuals in the sample. It is an approximation of the genome stability in a region and may be better at identifying unstable genomic regions than considering simpler statistics such as proportion of rearranged sites or number of SVs relative to a single reference. The reason we developed this statistic was because we aimed to better capture the potential evolutionary rate of a region. For example, if considering the fraction of non-syntenic bases, a single large variant would create a high value, even if it is the only variant present. On the contrary, many diverse, small SVs would possibly cause a deflated estimate of the SV diversity of the region if their total length was a small proportion of the region’s overall length. Though we do not present a detailed exposition of the method here, we present it as an intuitive and hopefully useful complementary technique for investigating structural diversity in pan-genomes. Our software for its calculation across sliding windows, svstats, is freely available on GitHub (https://github.com/markcharder/svstats). We used the program in this study to calculate SVpi in 50 Kb sliding windows across the genome with an increment of 1 Kb.

### Analysis of linkage disequilibrium and recombination

To assess linkage disequilibrium decay with physical distance, linkage disequilibrium was first calculated for all pairs of variants between variants with the plink flags ‘--ld-window-r2 0’, ‘–ld-window-kb 300’ and ‘--r2 dprime’. R^2^ was averaged for each physical distance and the distance at which average R^2^ reached half its maximum value was recorded. The program phipack (obtained from https://www.maths.otago.ac.nz/~dbryant/software/PhiPack.tar.gz) was used to conduct three tests of the association between distance and linkage disequilibrium, the pairwise homoplasy index, maximum Χ^2^, and nearest neighbour score tests.

To assess recombination rate, we selected four genotypically fairly uniform populations that had no obvious population structure. Recombination rate was calculated for these populations using ldhat version 2.2 [57] and recombination hotspots were identified with ldhot version 8.30 [106]. To run ldhat and ldhot, plink was first used to convert the pan- genome VCF file to plink PED and MAP files with the flags ‘--recode’, and ‘--biallelic-only’ and ‘--snps-only’ to keep only biallelic SNPs. These were used as input for the command ‘plink2ldhat’ from our program ‘svstats’ (https://github.com/markcharder/svstats) to convert to ldhat or ldhot format. A finite sites version of Watterson’s theta was calculated using the command ‘watfsites’ from svstats to provide a parameter for generating ldhat lookup tables. The ldhat program ‘complete’ was then used, with flags ‘-rhomax 100’ and ‘-n_pts 101’, and the appropriate number of ‘-n’ individuals, to create look-up tables for calculating variable recombination rates across chromosomes. The ldhat interval program was then used, with the appropriate look-up table, to calculate variable recombination rates with the flags ‘-exact’ ‘-its 10000000’ and ‘-samp 3500’. Reversible jump Monte Carlo Markov Chains were run starting with block penalties ranging from 5 to 50 (with an increment of 5) and chains were assessed for convergence. Posterior distributions of rates and bounds from the chains were estimated using the ldhat command ‘stat’, with the flag ‘-burn 35’. Using the output of ldhat interval, ldhot was run using the appropriate look-up table with the additional flag ‘--nsim 1000’.

### Assessment of correlation between population-wide statistics and genomic features

The command ‘makewindows’ from Bedtools version 2.27.1 [107] was used to create sliding windows of 50,000 bp, with an increment of 1,000 bp, across the *S. sclerotiorum* genome. To calculate gene and repeat density, the bedtools command ‘coverage’ was used with Braker3 and EDTA annotations, respectively, and the sliding windows. To calculate GC content for windows, the command ‘nuc’ was used. Methylation rates from bisulfite sequencing data were converted to BED format using a custom script in R, and bedtools ‘intersect’ with the flag ‘-c’ was used to calculate the number of methylated sites per sliding window. A BED file was also created from the ldhat output, using a simple Awk script, and used to calculate recombination rate for sites in sliding windows. The rate was summed across sliding windows for comparison. Comparison between recombination rate and other statistics of interest was conducted in R using Spearman’s rank correlation.

The bedtools command ‘closest’ was used to determine the distance between transposon annotations from EDTA and the nearest structural variant in all genomes. A Kruskall-Wallis test in R was then used to determine whether any transposon classes were significantly closer than other classes to the nearest SV.

### Genome-wide association and trait correlation analyses

Before conducting GWAS and whole genome regression analyses, phenotype data were normalised with the R package bestNormalize version 3.5. Two GWASs were run. One (GWAS1) used variants that were filtered so that they were in approximate linkage equilibrium using plink version 1.9 with the flag ‘--indep-pairwise 50kb 50 0.8’, whereas the other (GWAS2) did not. Both GWASs were conducted using GAPIT [108] with the BLINK model. This model has been shown to adequately correct for population structure whilst maintaining statistical power, and there were no non-genetic confounding factors between populations as phenotypic data were collected in the same environment. We therefore included no further population structure correction with, for example, principal components or a kinship matrix. GWAS1 was used to identify significant marker trait associations as it had fewer correlated markers than GWAS2 and therefore more statistical power. GWAS2 was used for the comparison of average absolute effects from structural and non-structural variants.

To determine whether SVs had a larger impact on traits than other variants, we conducted three tests. Firstly, we simply used standard t tests to compare the mean distributions of absolute effect sizes of non-SVs and SVs. Since SVs had a lower minor allele frequency on average than non-SVs, and this could affect variance of the test statistic, we developed a randomisation test. This test sampled non-SVs 500 times, each time creating a random set of non-SVs matching in number the total count of SVs. This random set was sampled so that proportions of variants with all possible minor allele frequencies (rounded to three significant digits) matched the minor allele frequency proportions in the SV set. The average absolute effect size from GWAS2 was recorded for each of these 500 samples and the number of times this effect size was larger than or equal to that of the mean absolute effect size of the SVs was treated as the empirical P value.

In our third test, we partitioned variants into those that were in linkage disequilibrium with structural variants and those that were not. We did this by first creating a file recording R^2^ for all pairs of neighbouring variants within 2 Kb with the plink flags ‘--r2’ and ‘--ld-window- kb’. From this file, we created a list of variants that had an r2 of >= 0.5 with at least one SV. This list, combined with the list of SVs themselves, was used to create two VCF files, one containing SVs and variants in approximate linkage disequilibrium with them and the other containing variants that were not SVs and were not in linkage disequilibrium with any SVs. The two VCFs were filtered so that variants were not in strong linkage disequilibrium with the plink command ‘--indep-pairwise 50kb 50 0.8’. Genomic relationship matrices [109] were created for each of these sets of variants and for the whole set of variants used in GWAS1 with the plink flag ‘--make-rel square’.

To assess genetic correlations between traits and determine whether adding SVs improved predictive ability, we fit univariate and multivariate linear mixed models with the R package sommer version 4.3.4 [110]. In these models, random effects for individuals were estimated with assumed variance and covariance described by the genomic relationship matrix proposed by Yang et al. (2011) [109]. To assess trait genetic correlations, the genomic relationship matrix was estimated using all biallelic variants of minor allele frequency >= 0.05. To assess improvement of prediction accuracy when including SVs, we fit models with one random effect with variance structured by a relationship matrix estimated with only non-SVs, and models with two random effects, one structured by a non-SV and the other structured by an SV-only relationship matrix. For each trait, we performed ‘leave-one-out’ cross validation. For each recording of each phenotype for each strain, the recording was masked and the two models fit with this recording missing. The predicted BLUP value based on the rest of the strains was recorded for this version of the model and Pearson’s correlation coefficient between model predictions and phenotypes was recorded.

## Declarations

### Ethics approval and consent to participate

Not applicable

### Consent for publication

Not applicable

### Availability of data and materials

The datasets generated and/or analysed during the current study are available in NCBI, under BioProjects PRJNA1112094 and PRJNA1120954, in the supplementary material, or available upon reasonable request from the corresponding author.

### Authors’ contributions

MCD collected data, designed and executed experiments, oversaw research, conducted all main analyses and wrote the first manuscript draft. YK designed and executed experiments and edited the manuscript. TN designed and executed experiments and edited the manuscript. PM collected data, oversaw research, and edited the manuscript. SJB oversaw research and edited the manuscript. ARL collected data. CG-T collected data. SC designed and executed experiments and edited the manuscript. LB collected data and provided feedback on the manuscript. C Camplone executed experiments. SV executed experiments. DH designed and oversaw experiments and edited the manuscript. C Coutou designed and executed experiments and edited the manuscript. LGK initiated the project, oversaw research and edited the manuscript. All authors read and approved the final manuscript.

## Acknowledgements

This work was supported by resources provided by the Pawsey Supercomputing Research Centre’s Setonix Supercomputer (https://doi.org/10.48569/18sb-8s43) and Nimbus research cloud (https://doi.org/10.48569/v0j3-qd51), with funding from the Australian Government and the Government of Western Australia.

## Supplementary Material

### Supplementary Figures

**Supplementary Figure 1. A dendrogram showing the percentage of alleles identical by state between strains in the collection.** The green vertical line shows the cutoff used to identify groups of individuals representing a single clone (blue).

**Supplementary Figure 2. The relationship between recombination rate (y axis) and coding sequence density (x axis) of 50 Kb sliding windows.** The line is a general additive model and the shading represents 95 % confidence intervals.

**Supplementary Figure 3. SVπ and repeat content in 50 Kb windows across the genome.** The same as Figure 3 D but shown for all chromosomes.

**Supplementary Figure 4. Q-Q plots for GWASs conducted for all traits.** The y axis shows observed P values and the x axis shows the expected P values given a normal distribution. All plots show that most points are on (adequate correction) or below (over-correction in some cases) the line, and P values are not inflated.

### Supplementary Tables

**Supplementary Table 1. A** BUSCO scores for all strains used to construct the pan-genome graph. **B.** Gaps and telomeres in each chromosome of each assembly. In the TELOMERES column, L stands for ‘left’ and R stands for ‘right’, referring to the two (arbitrary) ends of the chromosome in the assembly FASTA.

**Supplementary Table 2.** Strains, excluding the reference strain, 1980, used to create the *Sclerotinia sclerotiorum* pan-genome and call structural variants. Strains with Nanopore and Illumina data were used to create the pan-genome graph whereas strains with only Illumina data (previous or current study) were used for mapping and variant calling against the graph.

**Supplementary Table 3.** Results of phipack tests for recombination across the 120 independent *Sclerotinia sclerotiorum* lineages. These include the Neighbour Similarity Score (NSS), the Maximum Chi^2 (MAX_CHI2), and the Pairwise Homoplasy Index (PHI) tests. All tests were significant, with a P value of zero, indicating increasing levels of recombination between alleles with distance.

**Supplementary Table 4.** Recombination hotspots identified relative to the *Sclerotinia sclerotiorum* reference genome. Four non-structured population subsamples were used to identify hotspots.

**Supplementary Table 5.** Cytosine methylation data from alignment of bisulfite sequencing reads to the 1980 genome. The first column is the NCBI chromosome accession. The columns for these tables are described in the CLC genomic workbench manual here: https://resources.qiagenbioinformatics.com/manuals/clcgenomicsworkbench/current/index.php?manual=Call_Methylation_Levels.html. Each spreadsheet represents one of the samples, for example, 0 HPI R1 is 0 hours post-inoculation replicate 1.

**Supplementary Table 6.** Results of a Kruskal-Wallis test to determine whether LTR transposons were significantly closer to structural variants across all genomes than other transposons. Transposon classifications are taken from EDTA.

**Supplementary Table 7.** Transposable element content of the 24 Sclerotinia sclerotiorum genomes based on EDTA annotations.

**Supplementary Table 8.** Spearman’s correlation between estimated recombination rate and SVπ, SV count and transposon content of 50,000 bp sliding windows. Rows coloured in green are significant positive correlations, those in red are significant negative correlations and those not coloured are not significant. Overall, the majority of chromosomes and populations showed a correlation between recombination rate and both SVπ and SV count but not transposon content.

**Supplementary Table 9.** Functional terms associated with genes in the largest gene bubble. Results are from an InterProScan analysis. The Gene IDs are based on a cd-hit grouping of Braker3 annotations across all genomes.

**Supplementary Table 10.** Genetic and actual correlations between life history traits. Where genetic correlations are above 1, below -1 or ‘NA’, the model was likely poorly or over-fit.

**Supplementary Table 11.** Tests for overall impact of SVs on phenotype. Grey cells are for test statistics that were not significant. Green cells are for test statistics that indicate in increase in SV impact on phenotype. Red cells are for test statistics that indicate a decrease in SV impact on phenotype.

**Supplementary Table 12.** Linear mixed models testing improvement in predictive ability (Pearson’s ρ) from models with no SVs in the genomic relationship to matrix to models with two terms, one for SVs and the other for non-SVs, or to models with only SVs. Improvements in predictive ability were variable but some traits showed a relatively large improvement.

**Supplementary Table 13 A**. Results of a GWAS for 14 life history traits. **B** BLASTp hits for gene downstream of 48 bp InDel azoxystrobin QTL, which encodes a centrosomin. **C** BLASTp hits for gene upstream of 48 bp InDel azoxystrobin QTL, which encodes a protein with no known functional domains.

**Supplementary table 14 A**. RNA sequencing data used for Braker3 annotation of genomes. **B** The host genomes used for filtering RNA sequencing reads used in Braker3 annotation.

