## Supplementary figures and images for "Cryptic recombination and transposition drive structural variation to shape genomic plasticity and life history traits in a host generalist fungal plant pathogen"

### Supplementary_Figure_1.png

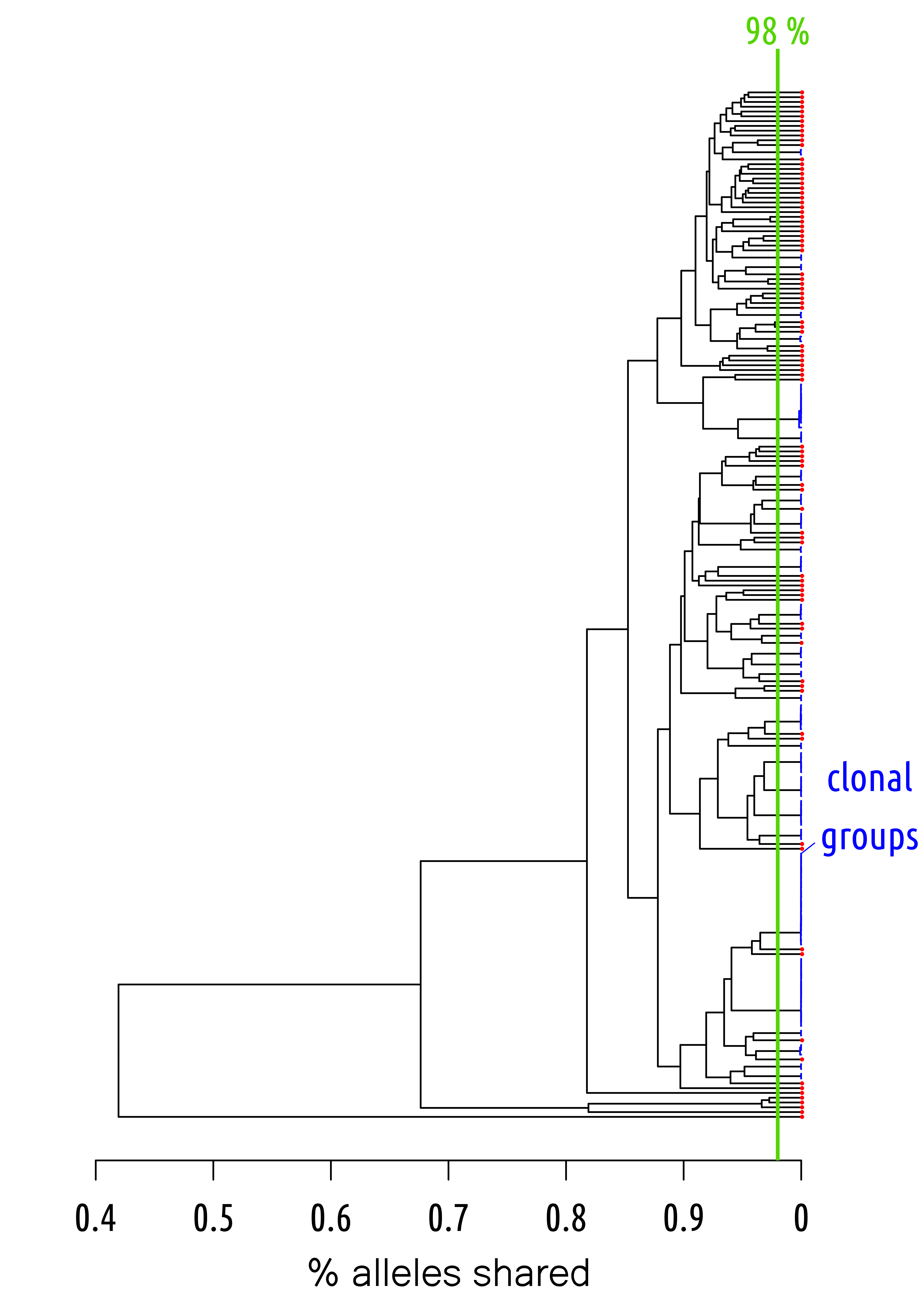

### Supplementary_Figure_2.png

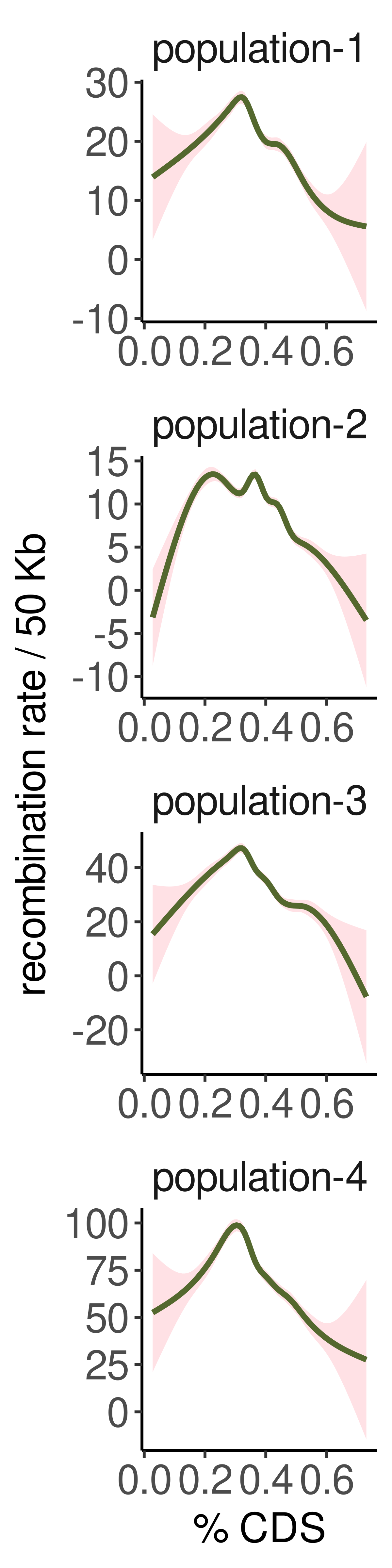

### Supplementary_Figure_3.png

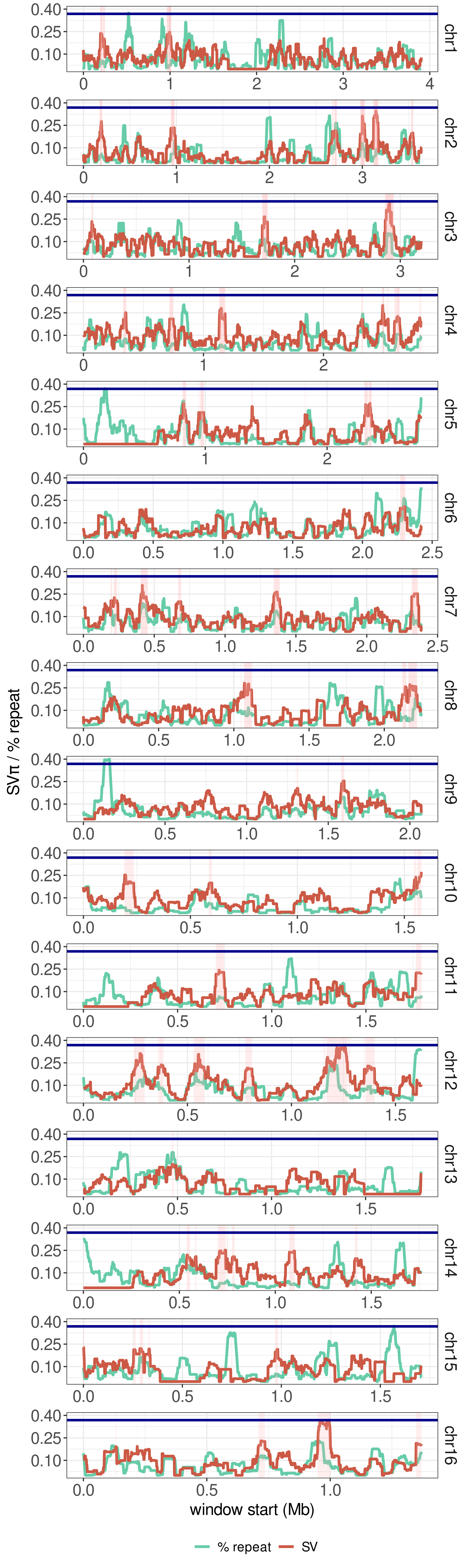

### Supplementary_Figure_4.png

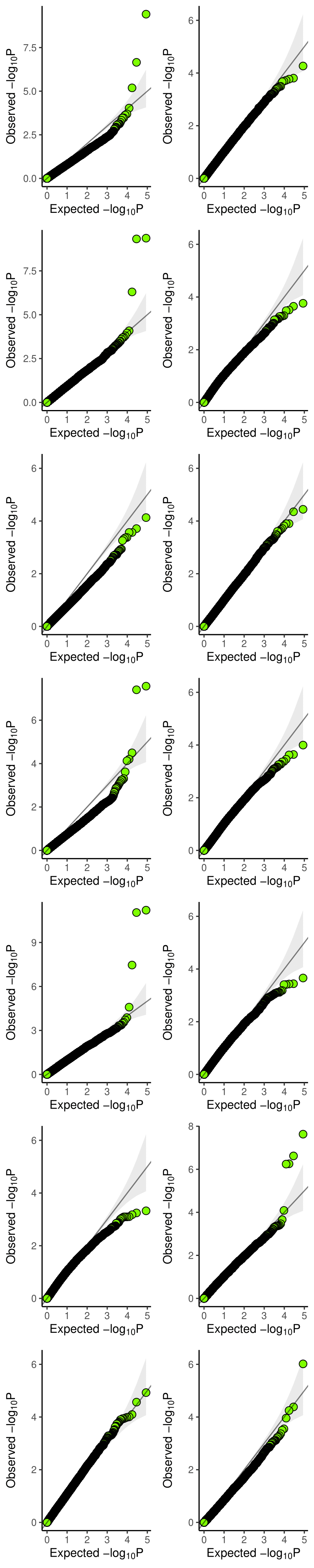
